# The haplotype-resolved genome sequence of hexaploid *Ipomoea batatas* reveals its evolutionary history

**DOI:** 10.1101/064428

**Authors:** Jun Yang, M-Hossein Moeinzadeh, Heiner Kuhl, Johannes Helmuth, Peng Xiao, Guiling Liu, Jianli Zheng, Zhe Sun, Weijuan Fan, Gaifang Deng, Hongxia Wang, Fenhong Hu, Alisdair R Fernie, Bernd Timmermann, Peng Zhang, Martin Vingron

**Author notes:** These authors contributed equally to this work (Jun Yang, M-Hossein Moeinzadeh). Correspondence should be addressed to P. Z. or M. V.

## Abstract

Although the sweet potato, *Ipomoea batatas*, is the seventh most important crop in the world and the fourth most significant in China, its genome has not yet been sequenced. The reason, at least in part, is that the genome has proven very difficult to assemble, being hexaploid and highly polymorphic; it has a presumptive composition of two B_1_ and four B_2_ component genomes (B_1_B_1_B_2_B_2_B_2_B_2_). By using a novel haplotyping method based on de novo genome assembly, however, we have produced a half haplotype-resolved genome from ∼267Gb of paired-end sequence reads amounting to roughly 60-fold coverage. By phylogenetic tree analysis of homologous chromosomes, it was possible to estimate the time of two whole genome duplication events as occurring about 525,000 and 341,000 years ago. Our analysis also identified many clusters of genes for specialized compounds biosynthesis in this genome. This half haplotype-resolved hexaploid genome represents the first successful attempt to investigate the complexity of chromosome sequence composition directly in a polyploid genome, using direct sequencing of the polyploid organism itself rather than of any of its simplified proxy relatives. Adaptation and application of our approach should provide higher resolution in future genomic structure investigations, especially for similarly complex genomes.

## Introduction

With a consistent global annual production of more than 100 million tons, as recorded between 1965 and 2014 (FAO), the sweet potato *Ipomoea batatas*, is an important source of calories, proteins, vitamins and minerals for humanity. It is the seventh most important crop in the world and the fourth most important crop of China. In periods of shortages of basic cereal foods, *Ipomoea batatas* frequently served as the main food source for many Chinese. It rescued millions of lives during and after three years of the Great Chinese Famine in the 1960s and was subsequently raised as a main guarantor of food security in China.

Although the sweet potato is such as outstanding crop, its genome has not yet been sequenced. The reason, at least in part, is that the genome has proven very difficult to assemble, being hexaploid (2n = 6x = 90) and highly polymorphic; with a base chromosome number of 15 and a genome size of about 4.4 Gb, as estimated from its measured C-value (Ozias-Akins and Jarret 1994). Sweet potato has a composition of two B_1_ and four B_2_ component genomes (B_1_B_1_B_2_B_2_B_2_B_2_), as predicted by genetic linkage studies using RAPD and AFLP markers (Ukoskit and Thompson 1997; Kriegner et al. 2003). The degree of homology, however, could not be estimated with accuracy since its genomic components are still largely poorly characterized. Recently, the genome survey sequencing of *Ipomoea trifida*, the most probable diploid wild relative of *Ipomoea batatas*, has been reported. Unfortunately, the current version of the *Ipomoea trifida* genome cannot serve as a reference sequence for *Ipomoea batatas* because of the low N50 value of its assembly, which is ∼36 Kbp, and even more because of the high abundance of gaps in the assembly, estimated at more than 30 percent(Hirakawa et al. 2015). This example also points out the problems inherent in the usual circuitous tactics employed in polyploid genome sequencing projects, which always begin with simpler diploid relatives. For example, sequencing of the autotetraploid potato (*Solanum tuberosum*) employed a homozygous doubled-monoploid potato (Consortium 2011). Allotetraploid Upland cotton (*Gossypium hirsutumdi*, AADD)(Li et al. 2015) began with diploid *Gossypium raimondii* (DD)(Wang et al. 2012) and *Gossypium arboreum* (AA)(Li et al. 2014). Comparably, the sequencing of allohexaploid bread wheat (*Triticum aestivum*, AABBDD) was initiated by sequencing the genomes of *Triticum urartu* (AA)(Ling et al. 2013) and *Aegilops tauschii* (DD)(Jia et al. 2013), but is still struggling with the precise sequencing of isolated chromosome arms(Choulet et al. 2014; Consortium 2014). All in all, a more cost-effective strategy and one that promises more efficient output for the direct sequencing of complex polyploid genomes is needed, especially for plant scientists, since polyploidy is a frequent and naturally-occuring state in plants. Although the full genetic implications of polyploidization are still obscure, it is clear that this state provides an important pathway for plant evolution and specialization. For the plant genome investigator, however, it remains a major problem.

*De novo* assembly of polyploid genomes remains a critical unsolved technical problem. The high heterozygosity that comes from the presence of three to six or even more copies of the monoploid genome will always hinder the genome assembly process, even with state-of-the-art assembling tools such as Platanus(Kajitani et al. 2014), MaSuRCA(Zimin et al. 2013) and SOAPdenovo2(Luo et al. 2012). The reason is that genome assembly focuses on the vast majority of bases that are invariant across homologous chromosomes; since these invariant regions are intermittent along the whole chromosome, the result is fragmentation in polyploid assembly. To address this problem, one can separately sequence each chromosome using isolated chromosome arms as reported in the wheat genome project(Choulet et al. 2014; Consortium 2014). The chromosome isolation technique, however, is tedious and has a long way to go before it is routine. Nevertheless, this strategy has inspired bioinformaticians to develop new tools for assembling each pair of homologous chromosomes separately. In contrast, with genome assembly, the haplotyping process pays more attention to DNA sequence differences among homologous chromosomes. Haplotyping of the human individual genome itself, however, poses its own challenges and the methods are costly, time consuming, and labor intensive (Duitama et al. 2011; Cao et al. 2015; Snyder et al. 2015). Since the distance of the adjacent variants between paternal and maternal chromosomes in humans is normally in the kilobase range (Consortium 2005), it is beyond the capacity of current cost-effective sequencing platforms to cover at least two variant positions in most cases. Nevertheless, human haplotyping studies have already indicated that genome assembly and accurate haplotyping are tightly linked (Cao et al. 2015). Unfortunately, the computational phasing problem in polyploidy is considerably harder than for a diploid organism because in the polyploid case one cannot make inferences about the “other” haplotype once one has seen the first (Aguiar and Istrail 2013; Berger et al. 2014).

Our initial sequencing of the *Ipomoea batatas* genome revealed that the distance between adjacent polymorphic sites is roughly one tenth of the distance in the human genome. Based on previous statistics, there are approximately ∼14 million polymorphic sites in the estimated 700∼800 Mb monoploid genome of *Ipomoea batatas*. This means that, on average, one read (100∼150bp length) from Illumina sequencing will cover 2∼3 polymorphic sites. This density of such sites should permit phasing the *Ipomoea batatas* genome, employing cost-effective Illumina paired-end sequencing. While the high heterozygosity of *Ipomoea batatas* makes genome assembly more challenging, it simultaneously makes haplotyping easier.

Here we report the development of computational tools to derive a genomic sequence from a polyploid species and one that is phased to a large degree. The haplotype-resolved *de novo* assembly of the *Ipomoea batatas* genome was generated entirely based on Illumina sequencing data. The final ∼824 Mb assembly has a scaffold N50 of ∼86 kb (Table 1). The entire number of scaffolds was 43,797 and their lengths varied between 3,92 to 803,129 bp. Altogether, there were 2,708 scaffolds longer than 86 kb. In total, 67,627 gene models were extracted and 76 gene clusters were identified. Furthermore, this consensus genome was phased into six haplotypes in 665,680 regions. Via phylogenetic analysis of these haplotypes, a hypothesis of the origin of modern cultivated *Ipomoea batatas* could be proposed and examined. In addition, the analysis permitted the estimation of the times of two recent genome wide duplication events; these were placed at approximately 525,000 and 341,000 years ago.

**Table 1.**
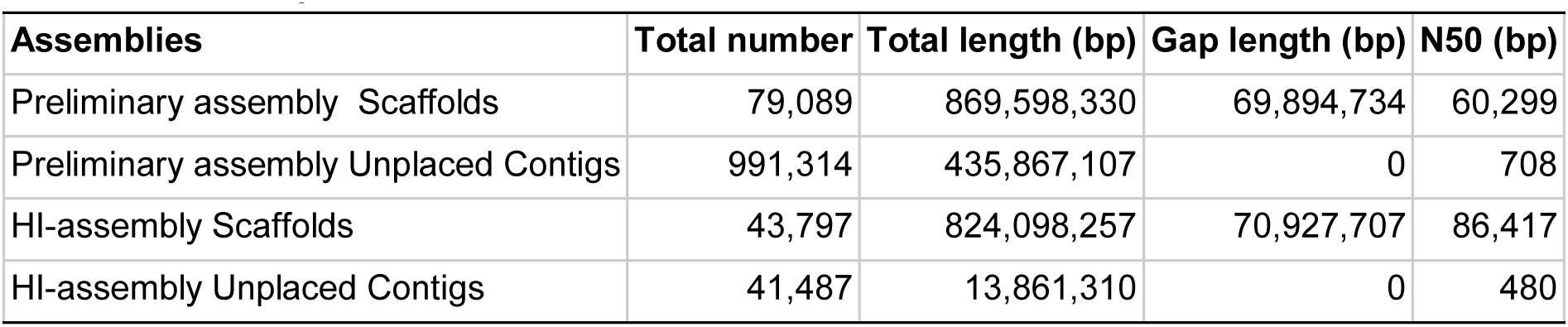
Summary of assemblies.

## Results

### Sequencing data generation

A newly bred carotenoid-rich cultivar of *Ipomoea batatas*, Taizhong6 (China national accession number 2013003), was used for genome sequencing (Figure 1). During the genome survey stage, three sequencing libraries were constructed and sequenced on Hiseq 2500 and GS FLX+ platforms (Supplementary Table 1, A500, A1kb and A454). After a preliminary genome assembly and read mapping for variant calling, the requirement for haplotyping of hexaploid *Ipomoea batatas* was estimated to be at least 40-fold monoploid genome coverage (Supplementary Figure 1). A new library was sequenced on the Nextseq 500 platform to meet the data requirement estimation (Supplementary Table 1, L500) and an additional gel-free mate pair library was also sequenced on the Nextseq 500 platform to improve scaffolding (Supplementary Table 1, MP). The insert size distributions of these paired-end libraries are shown in Supplementary Figure 2.

**Figure 1:**
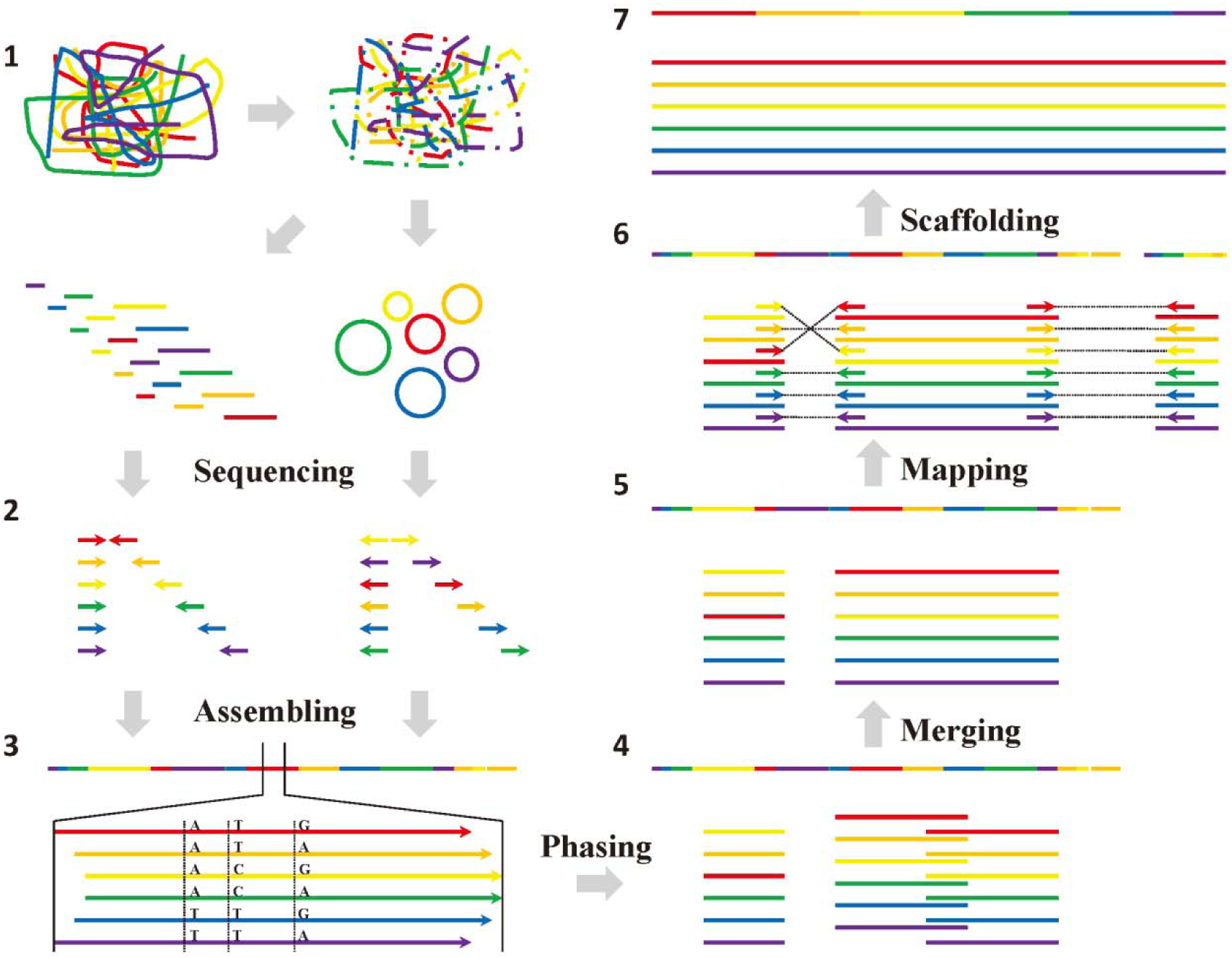
*De novo* assembly pipeline of haplotype-resolved hexaploid *Ipomoea batatas*. Genomic DNA was extracted from Taizhong6 *in vitro* plant and processed as follows: (1) DNA was fragmented into different sizes according to paired-end (lines) or mate-pair (circles) library requirements. (2) DNA sequences were obtained by Illumina sequencing of paired-end and mate-pair libraries. (3) *De novo* assembly of short reads and seed finding based on variant calling. (4) Phasing haplotypes by extending more strongly supported seed regions. (5) Merging overlapped haplotypes into longer haplotypes. (6) Mapping all raw reads against phased haplotypes. (7) Scaffolding based on haplotypes and consensus genome.

**Supplementary Table 1.** Statistics of QC-passed reads and mapped sequence data obtained from all libraries.

| Library | Platform | Sequencing type | Insert size | QC-passed reads | Mapped reads* | Mapped bases* (Gb) |
| --- | --- | --- | --- | --- | --- | --- |
| A500 | Hiseq 2500 | PE100 | 350 | 345,333,619 | 326,884,297 | 32.69 |
| A1kb | Hiseq 2500 | PE100 | 950 | 190,263,659 | 180,892,054 | 18.09 |
| L500 | Nextseq 500 | PE150 | 550 | 837,098,772 | 792,765,715 | 118.91 |
| MP | Nextseq 500 | PE150 | No size selection | 694,919,486 | 636,875,700 | 95.53 |
| A454 | GS FLX+ | SE (up to 1kb) | - | 3,385,694 | 3,348,732 | 1.68 |
| **Total** | - | - | - | **2,071,001,230** | **1,940,766,498** | **266.90** |

\* Map against preliminary assembly

**Supplementary Figure 1:**
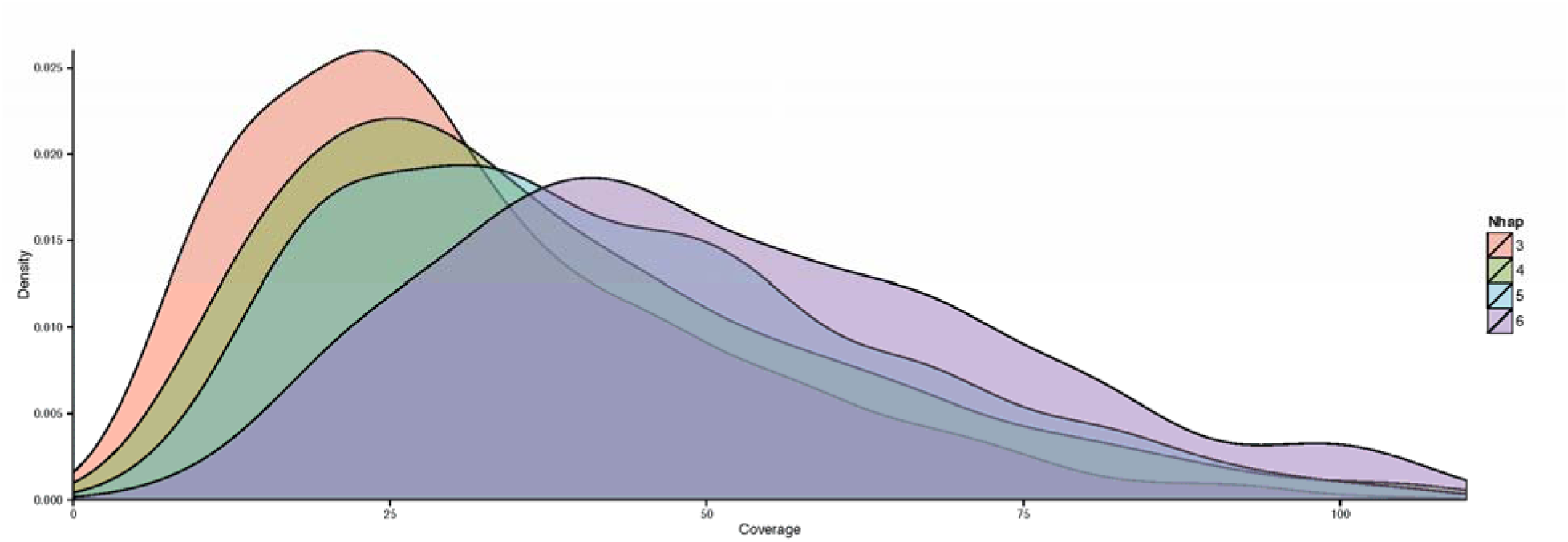
Coverage distribution of genomic regions phased into triploid, tetraploid, pentaploid, and hexaploid during genome survey. The peak coverage around 40 in hexaploid indicates the minimal sequencing depth requirement for haplotyping of the hexaploid genome. The peak coverage shifting from triploid to hexaploid demonstrates the insufficient sequencing depth in the genome survey stage.

**Supplementary Figure 2:**
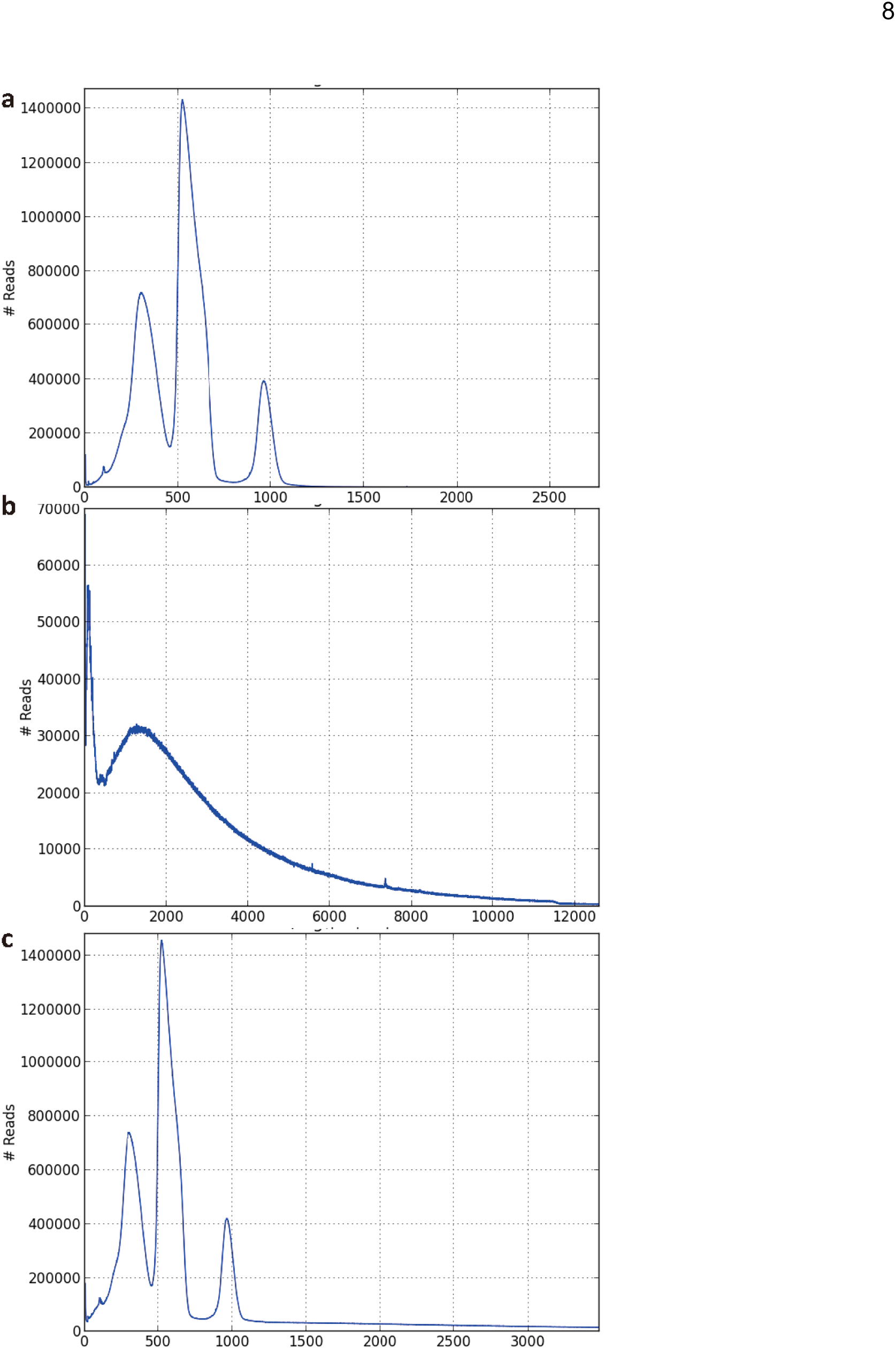
The insert size distribution of sequenced paired-end libraries. (a) Insert size distribution of libraries A500, L500, and A1kb. (b) Insert size distribution of mate pair library MP, truncated long tail at 12, 250 bp. (c) The Insert size distribution of all paired-end libraries, truncated long tail at 3,500 bp.

### Initial assembly of consensus genome

A heterozygosity-tolerant assembly pipeline, combining de Bruijn (Peng et al. 2012) and OLC (Margulies et al. 2005) graph strategies, was employed to carry out the hexaploid genome assembly of *Ipomoea batatas,* using error corrected Illumina reads (see Methods section). A total length of ∼870 Mb, mainly representing the monoploid genome, was assembled using this pipeline, with the largest scaffold being 3.7 Mb (corresponding to a completely assembled endophyte *Bacillus pumilus* genome). The second largest scaffold, which harbors 54 genes, was 581 kb (and contained 133 contigs). The N50 of all scaffolds was ∼60 kb with a 5,649 bp contig N50. The total number of scaffolds was 79,089 and their length varied between 312 and 3,723,026 bp. There were 3,796 scaffolds longer than 60 kb. Beside scaffolds, there were 991,314 contigs with a total length of ∼436 Mb. 92,790 contigs were longer than 1kb harboring a total of 175,679,534 bp. These contigs mainly reflected heterozygosity of the hexaploid genome since 97.35% contigs have been mapped back to scaffolds and one third of these contigs were found to match, despite many single nucleotide mismatches or small indels, as shown in Supplementary Figure 3. We call this version the “preliminary assembly”.

**Supplementary Figure 3:**
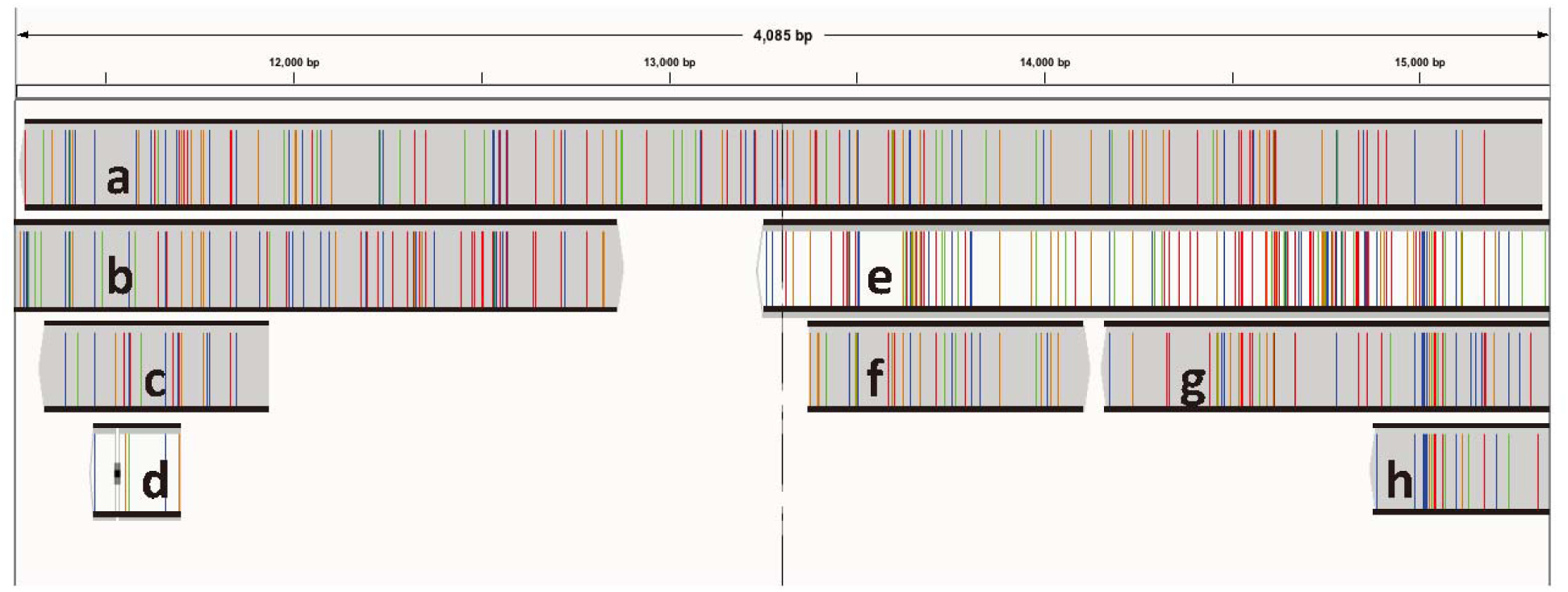
A snapshot of mapping results of contigs against scaffolds. The large number of mismatches indicates the single nucleotide polymorphism between homologous chromosomes in *Ipomoea batatas*. Cigar fields of these mapped scaffolds are listed as follows. (a) 4045M. (b) 33S44M1D136M1I10M6D149M15D21M1I1799M. (c) 579M. (d) 56M13D162M. (e) 15S2396M2I92M. (f) 733M. (g) 1260M. (h) 1014M1I162M1D136M. Among these, (b), (e), (g) & (h) are partially shown here.

### Variant calling

All the Illumina raw reads were mapped back to all scaffolds of preliminary assembly (Table 1). After removing the PCR duplications, there were 1,389,994,016 mapped reads in the final bam file for variant-calling using freebayes (Garrison and Marth 2012). In total, there were 14,090,421 variations, consisting mainly of single nucleotide polymorphisms but also indels, across the 869,598,330 bp assembly. Most of the variant positions harbored two possible alleles (Figure 2a). The distance between adjacent variations extended between 1 and 15,996 bp. These findings confirmed our earlier conclusion that the *Ipomoea batatas* genome is very heterozygous with, on average, one polymorphic site every ∼60 bp and a median distance of 21bp between polymorphic sites. The distance distribution peaked at 6 bp and only 8% (1,130,678/14,011,389) observed distances are longer than 150bp (Figure 2b). This forms the basis for phasing haplotypes using 100 to 150 bp Illumina reads. A high correlation (r = 0.975) between “Number of variations” and “Scaffold length” was found after excluding the endophyte genome (Supplementary Figure 4), which increases our chances for phasing the scaffolds.

**Figure 2:**
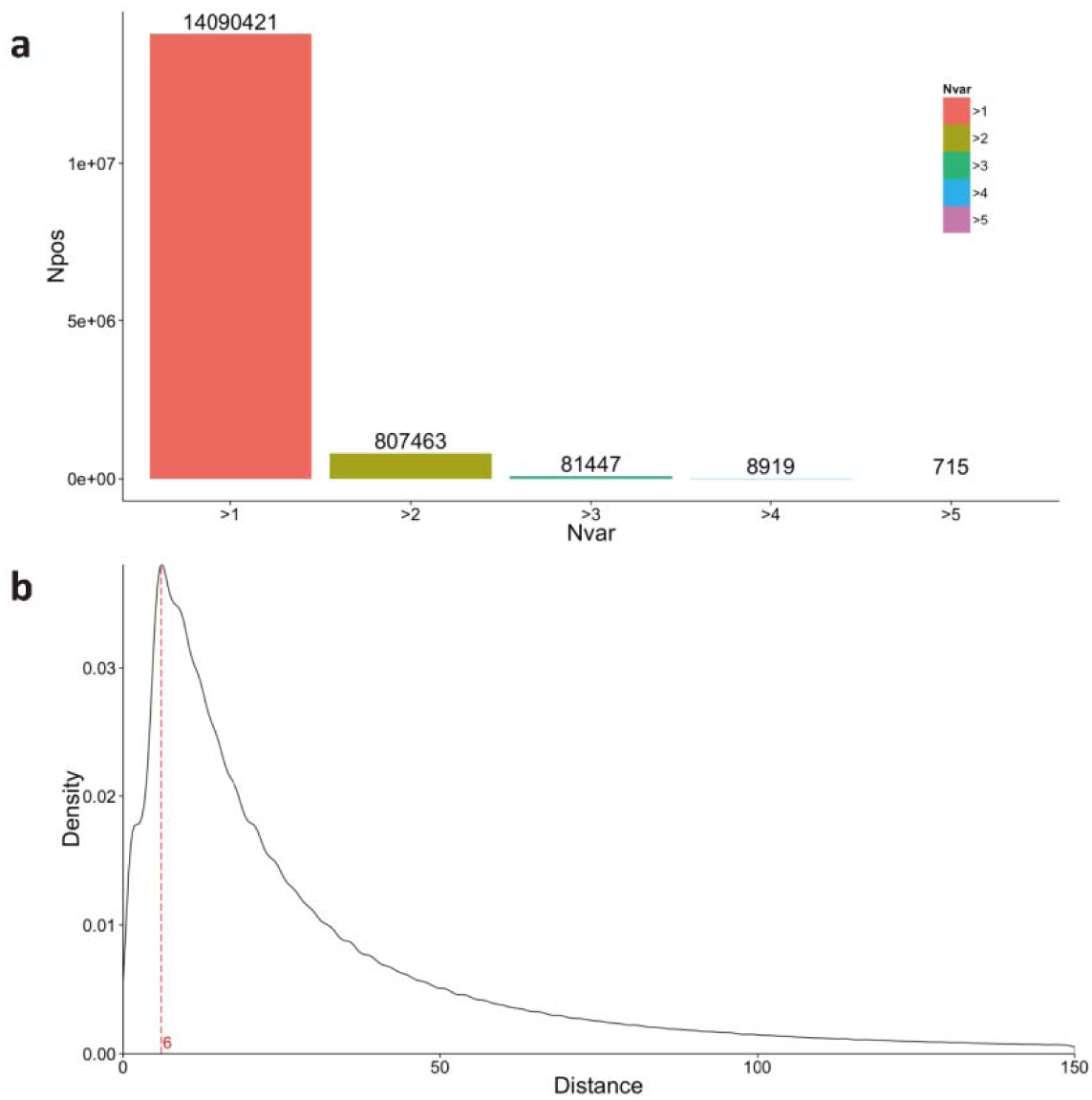
Summary of variations. (a) The total numbers of variation with different number of variant alleles. On average 869598330/14090421=61.7bp with one variant, which means short reads (100bp & 150bp) are informative for haplotyping. Npos, Number of positions; Nvar, Number of Variant. (b) Adjacent variation distance distribution peaked at 6 bp (red dashed line). Only 1130678/14011389=8% observed distances are longer than 150bp.

**Supplementary Figure 4:**
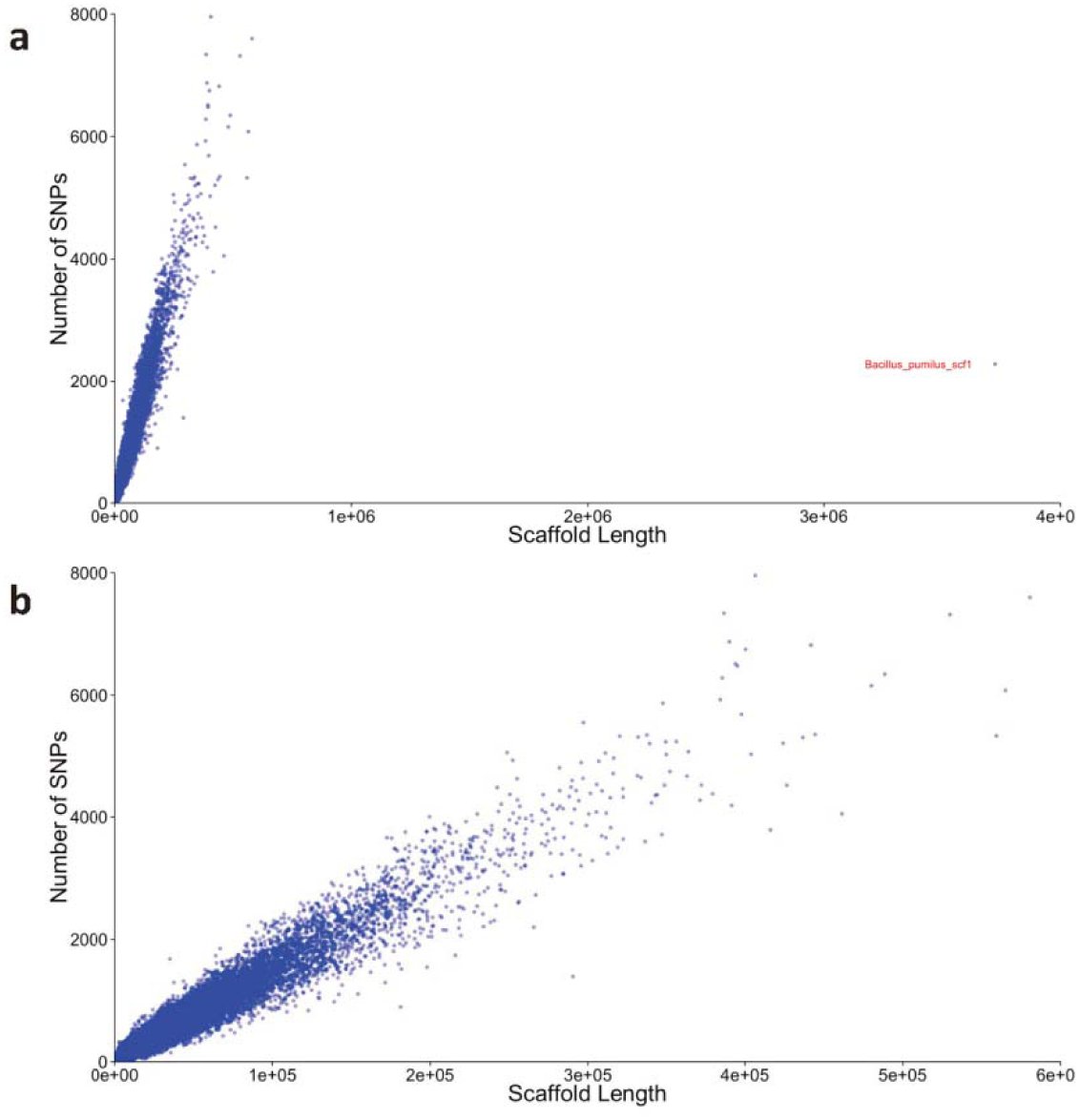
Summary of variations in scaffolds in preliminary assembly. (a) Number of variations and length of all scaffolds. An isolated dot represents the fully assembled genome of sweet potato endophyte, *Bacillus pumilus*. (b) High correlation (0.975) between “Number of variations” and “Scaffold length” after excluding the endophyte *Bacillus pumilus* genome.

### Phasing of haplotypes

For phasing, we developed an algorithm that assigns reads in a seed region of polymorphic sites to six presumptive haplotypes. For example, three polymorphic sites with two alleles each would give rise to eight combinations. Sequencing errors, however, could artificially inflate the number of hypothesized haplotypes. Thus, we looked for six haplotypes that have the most support in terms of sequencing reads (See method section and Figure 3). Haplotypes can be identified from combinations of alleles over many polymorphic sites that are connected in a read. Our algorithm looks for combinations of two, three, and four polymorphic sites. These phased regions were then extended continuously by searching for elongation of individual haplotypes. Finally, 602,450 regions were found and ∼40% of the genome was phased in to six haplotypes. These haplotypes could be further extended by connecting paired-end reads. Part of the paired-end reads link haplotypes within one assembly scaffold, while other paired-end reads connect haplotypes from different scaffolds which can then be used for “Haplotype-Improved assembly”.

**Figure 3:**
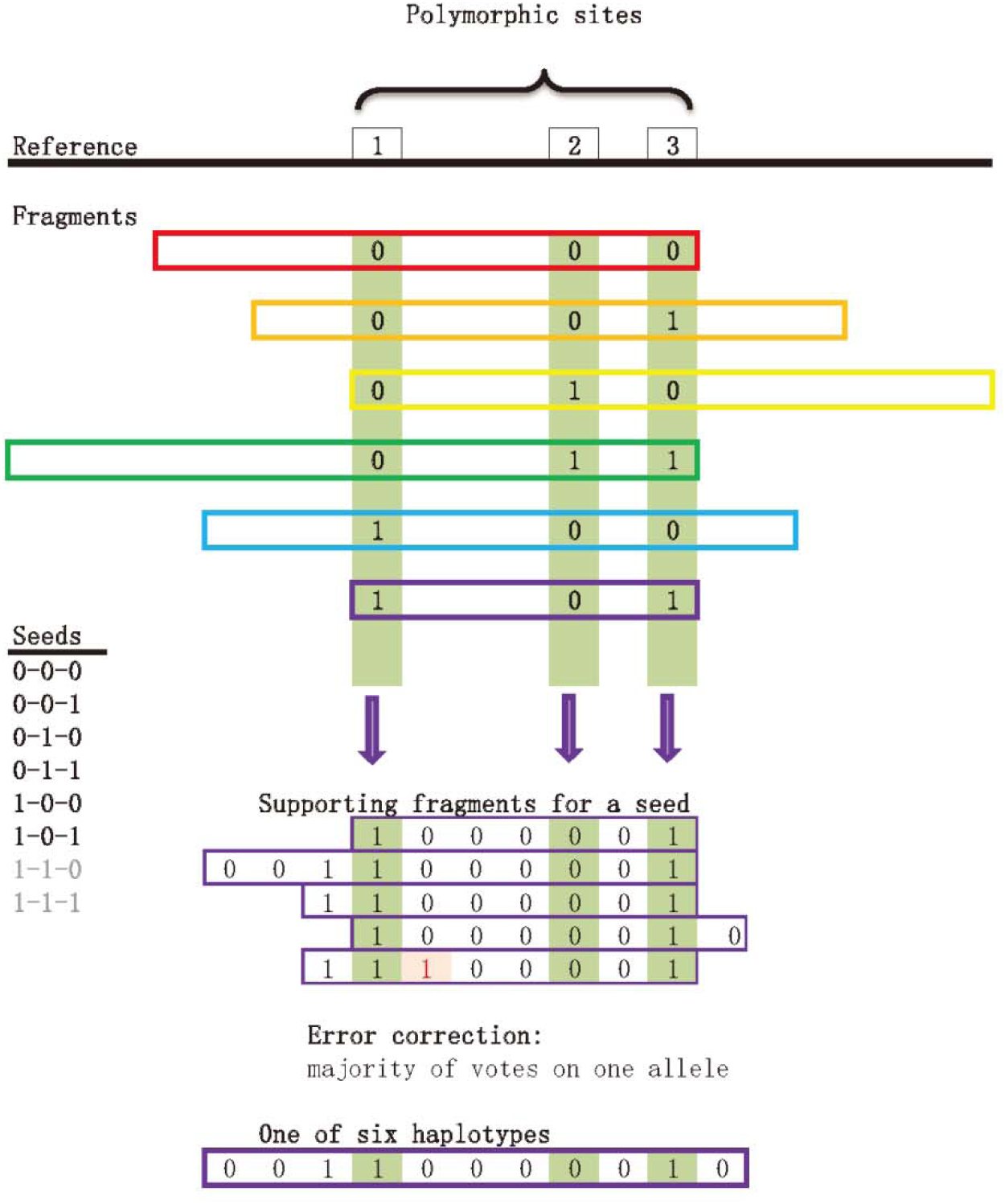
Illustration of seed finding algorithm. Step 1: Searching all possible seeds using different seed lengths. As an example, seed length 3 is shown here. Step 2: Selecting 6 highest supported seeds from all seeds found in previous step, gray color indicates less supported or hypothesized seeds. Step 3: Extension of particular seed based on supporting fragments. Reference: Only variant positions in each scaffold were extracted as reference and coded as 0. Fragment: Only variant positions in each read were extracted as fragment and coded by 0∼5 based on variation in particular position. Paired-end information was not used here. Seeds: Highest supported 6 seeds which can be used for distinguishing homologous chromosome regions. Haplotype: One haplotype represents a certain region in one chromosome.

### Haplotype-Improved assembly (HI-assembly)

All the Illumina raw reads were mapped against haplotype sequences generated by the phasing step. Only perfectly matched paired-end reads were considered as haplotype connections. The inter-scaffold and intra-scaffold connections were separated for haplotype-based scaffolding and haplotype elongation, respectively. We identified redundancy in the preliminary assembly based on exhaustive comparisons among scaffolds. If one scaffold was already covered by another longer scaffold with more than 85% sequence identity and more than 85% sequence length, the shorter one was removed. The largest scaffold in preliminary assembly, the endophyte *Bacillus pumilus* genome, was also excluded. After these procedures, there were 61,118 remaining scaffolds, of total length of 822,598,598 bp, with 64,561 bp N50. Then these scaffolds and haplotype-based connections were used as input files for the scaffolding software SSPACE (Boetzer et al. 2011). Gap sizes between connected scaffolds were estimated and super scaffolds were generated as the HI-assembly. The final assembly N50 of this consensus genome was ∼86 kb (Table 1). The total number of scaffolds was 43,797 and lengths varied between 392 to 803,129 bp. There were 2,708 scaffolds longer than 86 kb. When the contigs in preliminary assembly were mapped against the HI-assembly, one third of these contigs could be mapped back to full length though allowing for a fairly large number of mismatches. These contigs probably constitute particular haplotypes (Supplementary Figure 5). Using a similar scenario for removing redundancy in scaffolds, there were 41,487 remaining contigs, with a total length of ∼14 Mb (Table 1). The HI-assembly serves as our final assembly, which we annotate, use for final phasing of haplotypes, and from which we trace the evolution of the *Ipomoea batatas* genome.

**Supplementary Figure 5:**
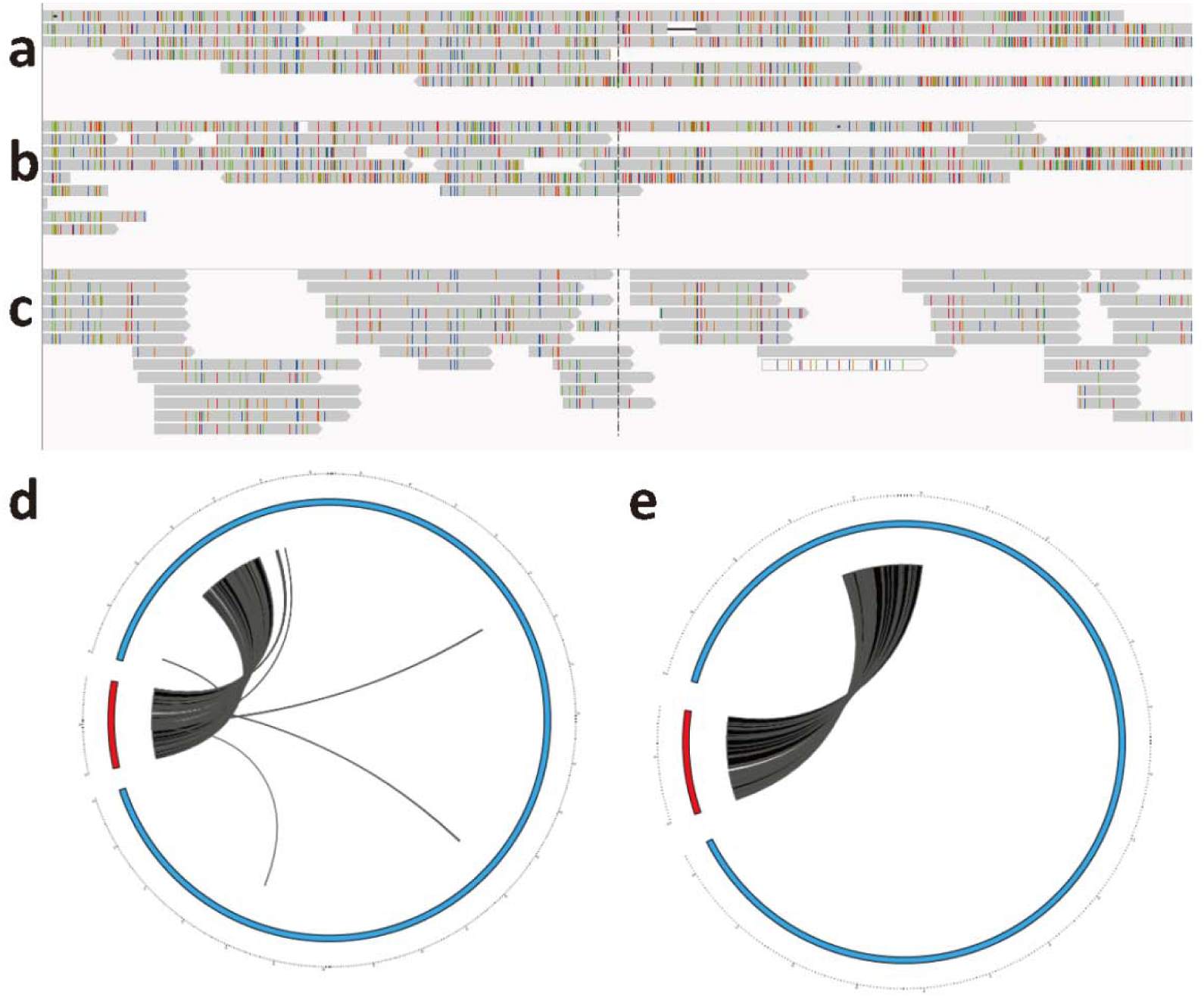
Comparison between scaffolds, contigs and haplotypes mapping results. (a) Scaffolds were mapped against scaffolds. (b) Contigs were mapped against scaffolds. (c) Haplotypes were mapped back to scaffolds. (d) Collinearity checking before removing of a large scaffold (19kb, red). (e) Collinearity re-checking before removing of a large scaffold (17kb, red). The higher density of mismatches in scaffolds and contigs (a and b) than haplotypes (c) indicates the problems of traditional assembling.

### Genome annotation and cDNA validation

There were 67,627 gene models extracted by StringTie (Pertea et al. 2015) after six transcriptome data sets (see methods) were mapped to the HI-assembly. To validate the predicted gene models from transcriptome data, 10,063 expressed sequence tags (ESTs) were generated by Sanger sequencing. These sequences were then mapped to the genome for validation of the predicted gene models. Of these, 9032 were located correctly and found to match the predicted gene models. We also mapped another 454 sequencing data set (Schafleitner et al. 2010) to the genome, and 95.69% of QC-passed reads were mapped and support the predicted genes. The functional annotations of predicted genes were obtained by homologous protein sequence searching in Uniprot (http://www.uniprot.org/) and Tair (https://www.arabidopsis.org/). The gene annotation and evaluation results are summarized in Table 2. RepeatModeler (Smit and Hubley, http://www.repeatmasker.org/RepeatModeler.html) was employed to identify repeat sequences in the present genome. The repeat sequence classifications are summarized in Table 3. The horizontally transferred T-DNAs reported by Kyndt, T. et al. were also investigated in the assembly (Kyndt et al. 2015). We found multiple copies of these T-DNAs suggesting that the horizontal gene transfer event has happened before hexaplodization of *Ipomoea batatas* (T-DNA1, NCBI Accession: KM052616, hits Scaffold171 and scaffold20897 and T-DNA2, NCBI Accession: KM052617, hits scaffold4674, scaffold4202 and scaffold121).

**Table 2.** Summary of gene annotation.

| Total number | Uniprot | Tair | EST* |
| --- | --- | --- | --- |
| 63,737 | 54,447 | 49,377 | 9,032 |

\* Gene model validation using sanger sequenced 10063 cDNA clones.

**Table 3.** Summary of repeat sequence identification.

| Type of elements | Number of elements | Length occupied | Percentage of genome* |
| --- | --- | --- | --- |
| LTR | 213,439 | 92,066,503 | 10.987 |
| DNA elements | 256,260 | 50,863,135 | 6.070 |
| LINE | 40,146 | 20,814,265 | 2.484 |
| Simple_repeat | 324,390 | 14,578,880 | 1.740 |
| RC/Helitron | 12,067 | 5,610,834 | 0.670 |
| Low_complexity | 45,912 | 2,324,352 | 0.277 |
| SINE | 4,168 | 618,723 | 0.074 |
| rRNA | 322 | 97,450 | 0.012 |
| Satellite | 456 | 33,443 | 0.004 |
| snRNA | 219 | 29,584 | 0.004 |
| Unknown | 985,978 | 195,232,724 | 23.299 |
| **Total** | **1,883,357** | **382,269,893** | **45.619** |

\* Scaffolds and Unplaced Contigs were taken as input sequences of RepeatModeler.

### Gene cluster identification

The discovery of operon-like gene clusters in plant genomes has prompted attempts to decode the regulatory mechanism in plant specialized compound biosynthesis (Nützmann and Osbourn 2015). Gene clusters of paralogous genes have been identified in a wide range of species including maize, lotus, cassava, sorghum, poppy, tomato, potato, rice, oat and Arabidopsis(Boycheva et al. 2014; Fernie and Tohge 2015). In the present case, the *Ipomoea batatas* genome, four 6∼7-gene clusters for alkaloids (Figure 4, GC002, GC006, GC014 & GC024), three 6∼8-gene clusters for terpenes (Figure 4, GC001, GC005 & GC031), and a 6-gene cluster for cellulose (Figure 4, GC028) were identified by searching orthologous genes using Exonerate (Slater and Birney 2005). In addition to the gene clusters shown in Figure 4, there were 68 more gene clusters found in the current genome assembly (Supplementary Table 2). The results indicate that pathway regulation via clustered genes is commonly used in *Ipomoea batatas*. Although all the orthologous genes found are based on protein sequence similarity, their biological functions are not necessarily the same as those reported in other species. Nevertheless, the identified gene clusters in *Ipomoea batatas* open up possibilities for investigating metabolic regulatory mechanisms in this plant.

**Figure 4:**
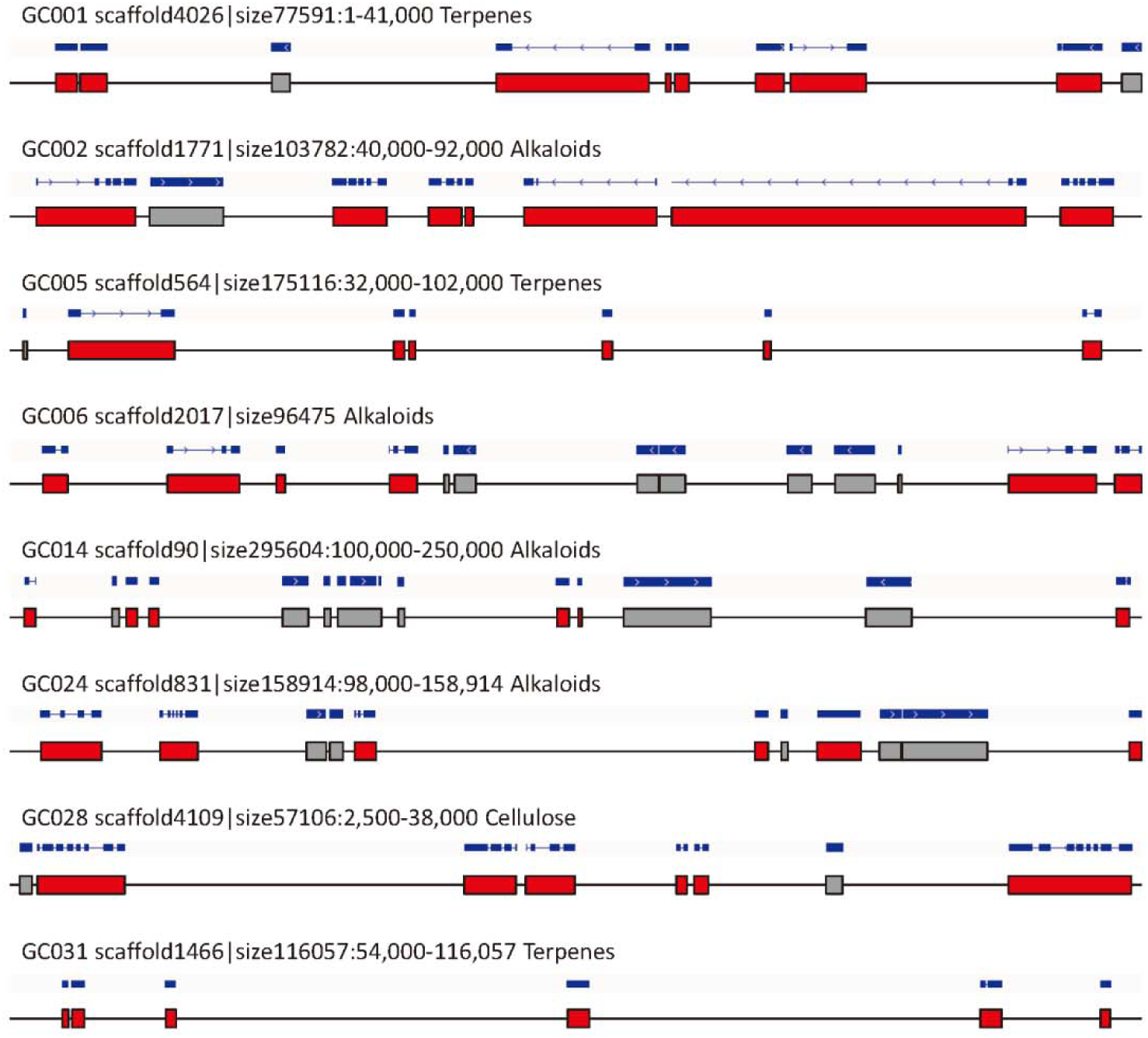
Identified gene clusters in present *Ipomoea batatas* genome. Blue bar panel indicated the gene structure, exon as box and intron as line. Red box indicated gene from the cluster and gray box indicated unrelated gene. The best hit genes of GC001 are *ZmBx4*, *AtMRO*, *AtMRO*, *PsCYP82X2*, *SbCYP71E*, *AtMRO*, *SlP450-1* and *SbCYP71E*; the best hit gene of GC002 is same, *SlGAME7*; the best hit genes for GC005 are *ZmBx5*, *PsCYP82Y1*, *OsCYP76M5*, *LjCYP71D11*, *PsCYP82X2* and *LjCYP736A2*; the best hit genes for GC006 are *PsCYP82X1*, *PsCYP82X2*, *OsCYP71Z6*, *ZmBx4*, *LjCYP79D4* and *PsCYP82X2*; the best hit genes for GC014 are *PsCXE1*, *PsCXE1*, *PsCXE1*, *SlGAME3*, *StSGT1* and *MeUGT85K4*; the best hit genes for GC024 are *SlGAME11*, *SlGAME7*, *SlGAME7*, *SlGAME3*, *SlGAME3* and *SlGAME3*; the best hit gene of GC028 is same, *StCeSy*; the best hit genes for GC031 are *MeCYP71E*, *OsCYP71Z7*, *OsCYP99A2*, *SlP450-1*, *OsCYP71Z6* and *SlP450-1*. All the detailed gene cluster information can be found in Supplementary Table 2.

**Supplementary Table 2.**
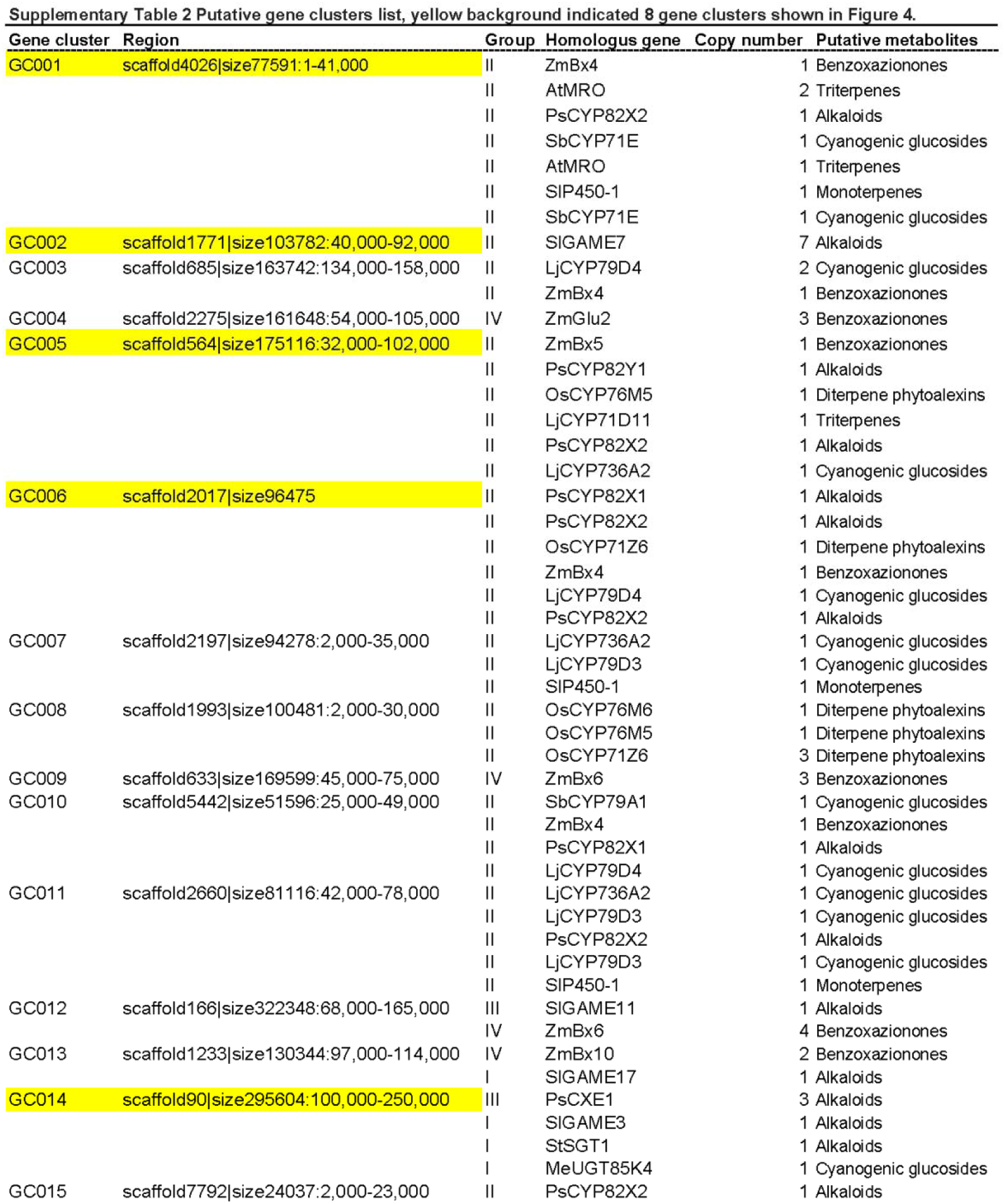
Putative gene clusters list (A long list with 182 rows is provided as an additional file).

### Updating and validation of haplotypes

Based on HI-assembly, we applied our phasing procedure a second time and phased 665,680 regions into 1,858,352,820 bp haplotype sequences. Thus, the new assembly leads to extended haplotypes in 10.5% more regions and a 12.9% improvement of total length. We validated these final haplotypes using a set of 454 reads (Supplementary Table 1, A454) that had been produced earlier but which had not been utilized for assembly or phasing. The reads are on the order of 1000 bp long and can thus serve to identify errors in the haplotype reconstruction. A large fraction of these 454 reads display haplotypes that have been correctly reconstructed by our short-read based methodology. More than 60% of overlaps between haplotypes and 454 reads are identical at variant loci. The longest reconstructed haplotype contained 88 polymorphic sites without any mismatch indicated by 454 reads. There may be many longer perfectly reconstructed haplotypes that remain undetected due to the limitation of the 454 read length (Supplementary Figure 6).

**Supplementary Figure 6:**
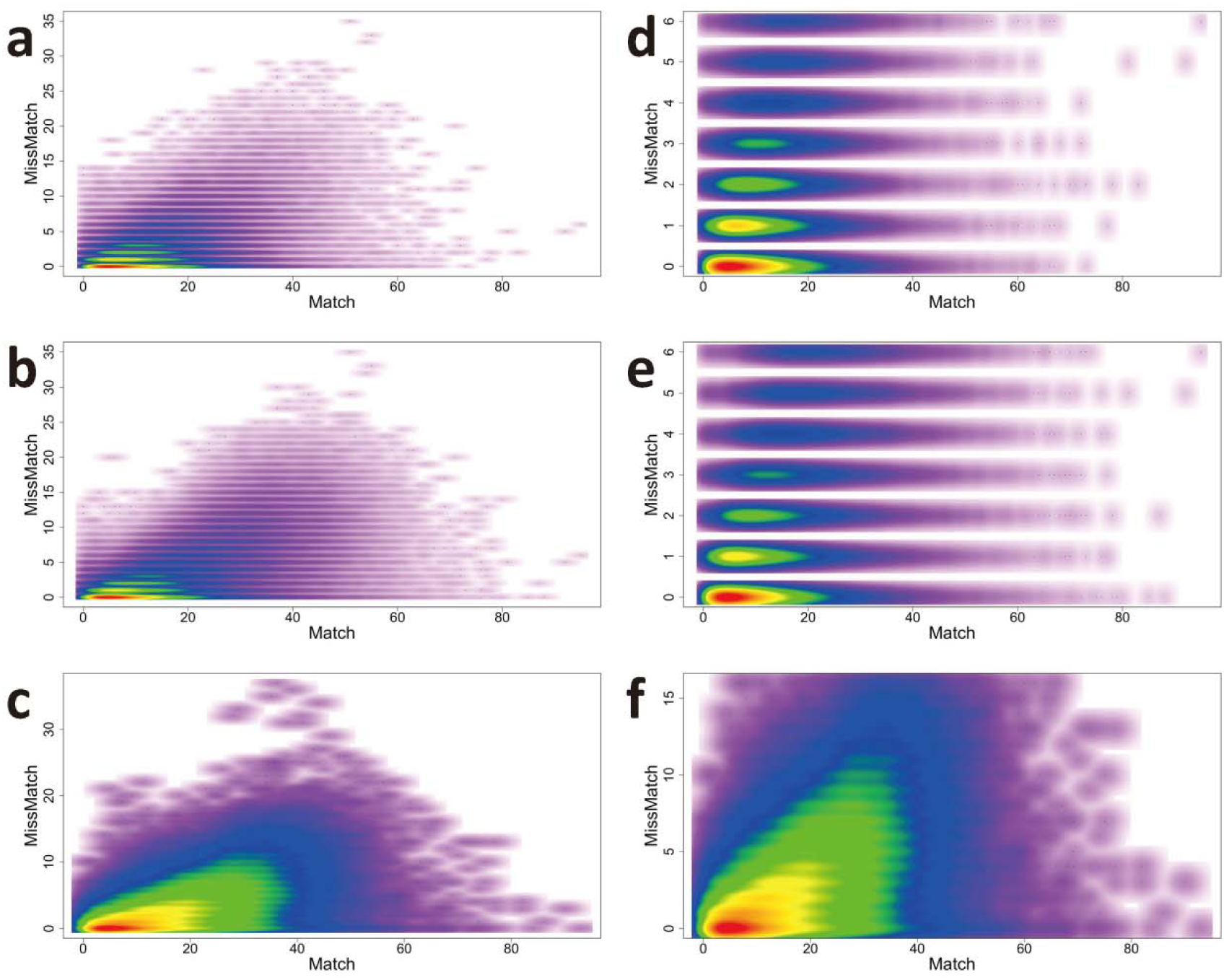
Haplotype evaluation by 454 reads. x axis: “match” is the number of coinciding polymorphic sites between haplotype and 454 read. y axis: “mismatch” is the number of different polymorphic sites in the overlap. Color indicates frequency of the respective (haplotype, 454 read) pairs, ranging from red to purple (on an exponential scale). (a) Evaluation after extension step. (b) Evaluation after bridge step. (c) Evaluation after paired-end connection step. (d) Less than 6 mismatch part of extension step (97.96% of total number of overlaps). (e) Less than 6 mismatch part of bridge (95.87%). (f) Less than 16 mismatch part of paired-end connection (99.31%). More than 60% of overlaps between haplotypes and 454 reads are identical at variant locus (d and e, y = 0). Even with strict mismatch threshold, a large fraction of 454 reads are supporting haplotypes reconstructed by short reads.

### Phylogenetic analysis of homologous chromosomal regions

Phylogenetic analysis was applied to the final haplotype-resolved genome. All the regions phased into six haplotypes were used for constructing UPGMA (Unweighted Pair Group Method with Arithmetic Mean) trees with the MEGA-Computing Core (Kumar et al. 2012). We generated 665,680 phylogenetic trees and determined their distribution over the six possible tree topologies explaining the evolution of the six haplotypes. The topology grouping of two haplotypes versus four haplotypes is the majority class. Furthermore, among the 4-haplotypes subgroup, 2 versus 2 dominates (Supplementary Figure 7). The average branch lengths of these 2:4-2:2 trees were obtained and a consensus tree was constructed (Figure 5a). These results suggest two whole genome duplication (WGD) events in *Ipomoea batatas* history together with a further diversification of the B_2_ subgenome (Figure 5b). Assuming a mutation rate of 0.7% base pairs per million years (Ossowski et al. 2010), the tetraploid progenitor of *Ipomoea batatas* was produced by a first WGD event estimated at 525,000 years ago. The origin of modern cultivated *Ipomoea batatas* would then have been the result of an initial crossing between this tetraploid progenitor and a diploid progenitor, followed by a second WGD event occurring about 341,000 years ago. The most probable diploid progenitor of *Ipomoea batatas* is also the most likely ancestor of modern *Ipomoea trifida*, although the tetraploid progenitor is still unknown. It might be identified, however, by a genome survey of the genomes of modern tetraploid species in *Ipomoea* genus.

**Supplementary Figure 7:**
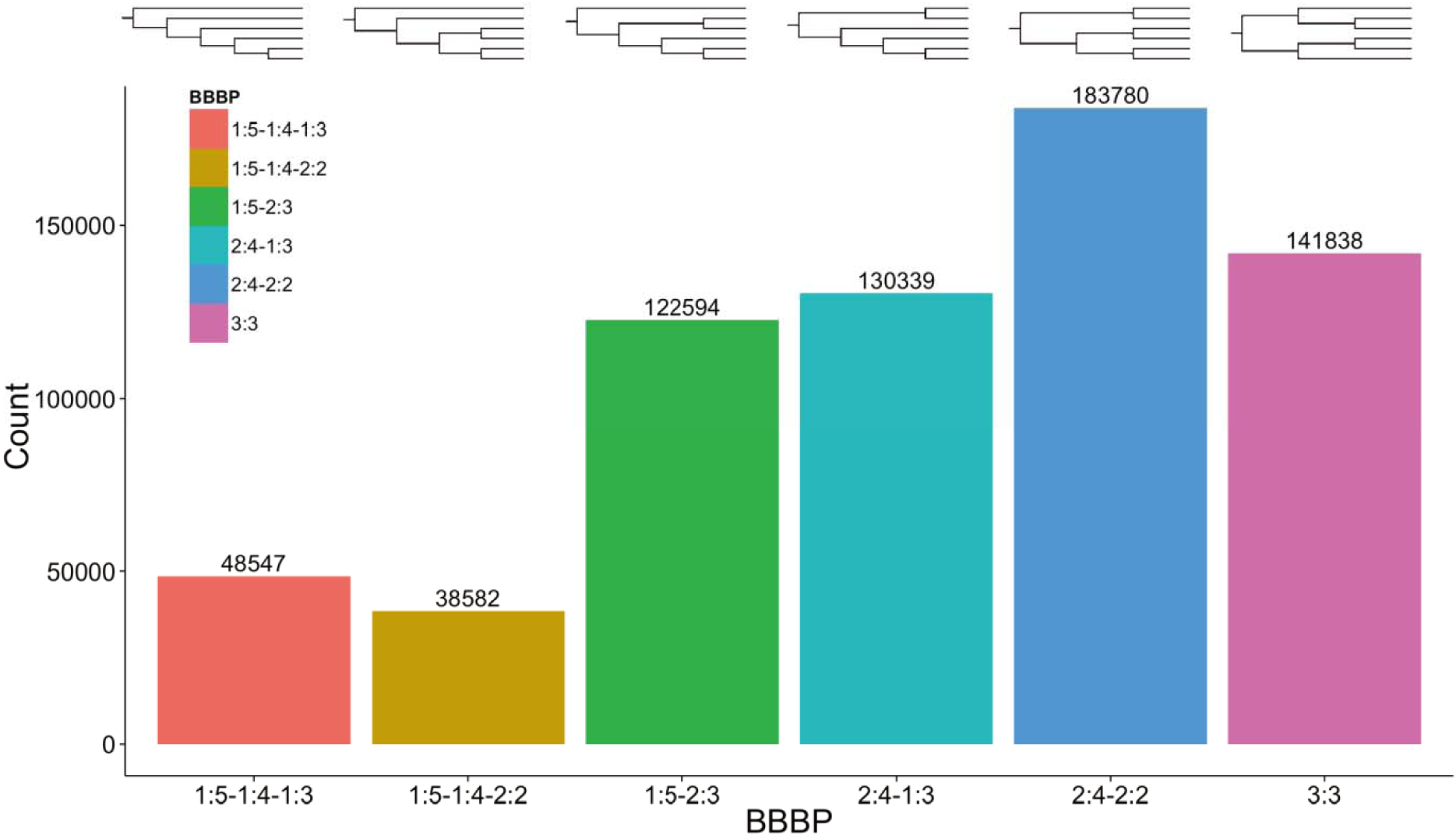
Distribution of six evolutionary topologies over all phased regions. The majority of the topologies for the 665,680 phased regions are grouped into 2 haplotypes versus 4 haplotypes (two blue bars). Among the 4 haplotypes subgroup, 2 versus 2 is dominant (light blue bar).

**Figure 5:**
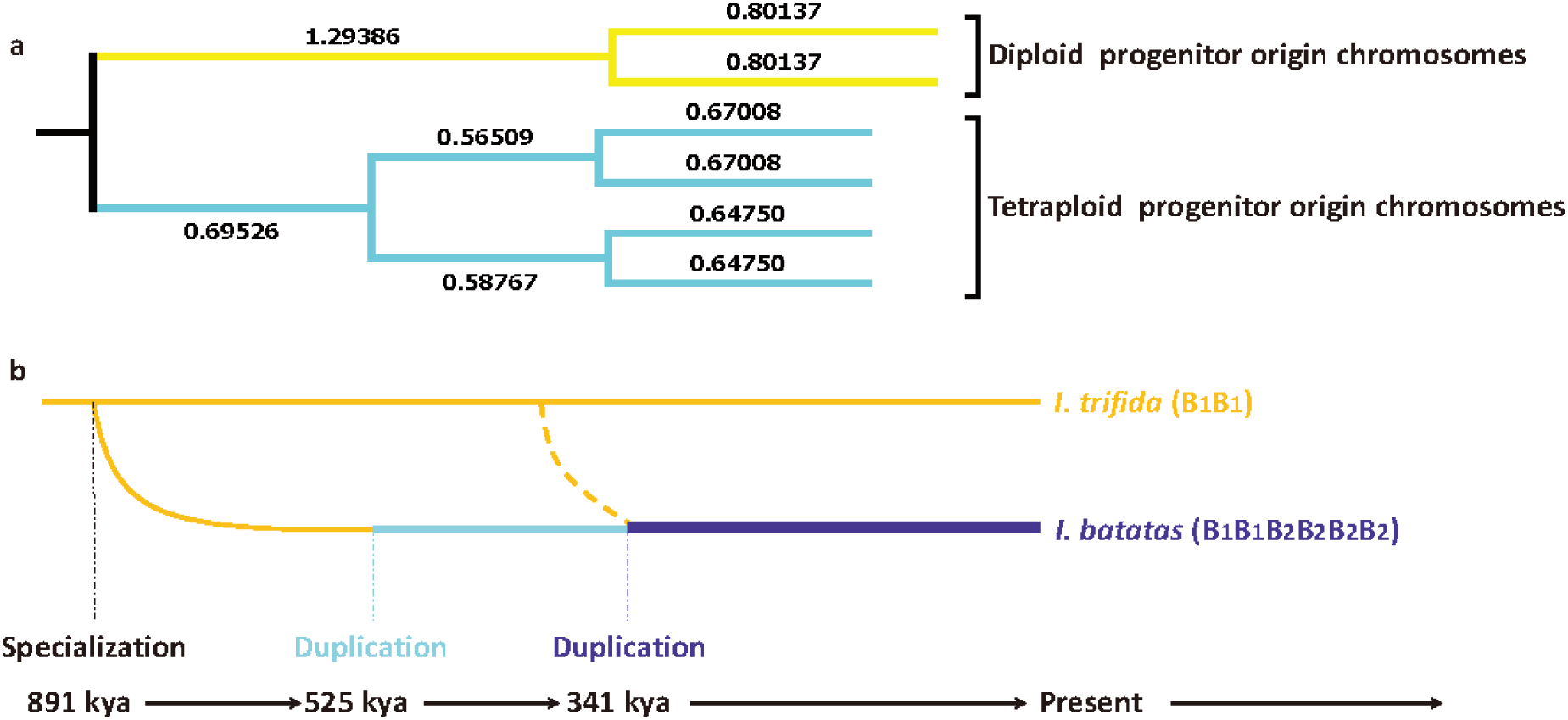
Evolutionary history of cultivated *Ipomoea batatas* revealed by phylogenetic analysis of homologous chromosome regions. (a) The dominant topology structure of all phylogenetic trees. Numbers indicated average branch length of trees in this structure. (b) The time points of B_2_ subgenome specialization and two whole genome duplication events were estimated as 891,000, 525,000 and 341,000 years ago. kya, a thousand years ago. Estimation based on 0.7% mutation rate per million years. Dashed curve indicated the crossing between diploid and tetraploid progenitors.

## Discussion

Here, we have reported the generation of sequencing data and the development of the necessary algorithms to compute long haplotype segments for the *Ipomoea batatas* genome. Based on an initial assembly and phasing, the computational process was repeated, yielding an improved assembly and extended haplotypes. In principle, haplotype phasing and haplotype-aided assembly can be iterated further to improve both results. In the present work, we only updated haplotypes based on the HI-assembly and observed remarkable improvements in terms of more phased regions and longer haplotype length. Optimization of the whole pipeline with iteration is now under way. Paired-end reads have been employed to extend haplotypes both within and across assembly scaffolds. Overall, high quality assembly and phasing can now be achieved with a comparatively low investment in sequencing (two runs on a HiSeq 2500 and two runs on a NextSeq; see Supplementary Table 1) but increased computational effort, both in terms of algorithm development and sheer computational power. Although the reconstruction of entire chromosomes remains out of reach by this methodology and is still reserved for alternative approaches including long-read sequencing technologies, many evolutionary questions regarding the ancestry of polyploid organisms can now be tackled using the methodologies we have developed. For the *Ipomoea batatas* genome, we have shown, based on the available haplotype alignments, that the majority of computed phylogenetic trees groups two haplotypes versus four, the latter being in turn symmetrically grouped into two and two. Although the different haplotype alignments obtained by our data and analysis do not allow identification of the full set of chromosomal connections among the segments, this phylogenetic pattern dominates. It is in agreement with a scenario in which modern cultivated *Ipomoea batatas* originated from a cross between a diploid progenitor and a tetraploid progenitor, followed by a whole genome duplication event. The precise timing of these evolutionary events, however, relies on an estimation of the average mutation rate for which we utilized the mutation rate reported for Arabidopsis (Ossowski et al. 2010). Nevertheless, we have demonstrated the power of phylogenetic analysis of the haplotypes derived by our approach in the reconstruction of the evolutionary history of *Ipomoea batatas.* Applied to the polyploidization of plants in general, phylogenetic analysis based on haplotype reconstruction could prove to be the most reliable way to study the origin of each set of chromosomes in complex polyploid genomes.

Our seed-based computational approach has thus proven successful on this very heterozygous genome, even with only short sequencing reads. This half haplotype-resolved hexaploid genome represents the first successful attempt to investigate the complexity of chromosome sequence composition directly in a polyploid genome, using direct sequencing of the polyploid organism itself rather than of any of its simplified proxy relatives. Adaptation and application of our approach should provide higher resolution in future genomic structure investigations, especially for similarly complex genomes. The pipeline presented here has a high degree of flexibility, which can be employed with many kinds of sequencing technologies and is almost ready to use for haplotyping tasks in a wide range of research programs. We are, of course, working on further improvements. With the availability of longer reads in the future, the same computational philosophy should be applicable for phasing of genome segments in other polyploid organisms, even when the density of polymorphic sites is lower.

## Methods

### Genome sequencing and assembly

A newly bred carotenoid rich cultivar of sweet potato (*Ipomoea batatas*), Taizhong6, whose China national accession number is 2013003, was selected as the target cultivar. Total genomic DNA was isolated from *in vitro* cultured plants as reported method (Kim and Hamada 2005). In total, five sequencing libraries were constructed and sequenced on Hiseq 2500, Nextseq 500 and GS FLX+ platforms (Supplementary Table 1, A500, A1kb, L500, MP and A454) according to the manufacturer’s instructions (Illumina, Inc. and Roche Applied Science), respectively.

The main steps of consensus genome assembly were as follows:

a. Read-correction of all Illumina data using the BFC package (https://github.com/lh3/bfc).
b. Assembling all short reads by IDBA-UD (with high kmers 100,123,150).
c. Further assembling the IDBA outputs by a long read assembler (NEWBLER 3.0).
d. Two scaffolding runs using the PLATANUS scaffolder on NEWBLER 3.0 output. During the first run, the median insert size of the libraries was set to the values observed in the distribution peaks (Supplementary Figure 2). In the second run, all the scaffolds from the first run were rescaffolded with a library median insert size of 6000 allowing huge standard deviations (6000±6000). It will therefore connect scaffolds in previous step using information from longer mate pairs presented in the MP library.
e. Gap-closing using all corrected illumina reads (PLATANUS GapCloser).

The detail information of the pipeline presented in this paper can be found in Supplementary Note.

### Haplotyping algorithm and pipeline

All the Illumina raw reads were mapped back to all scaffolds using BWA (Version 0.7.12-r1039). PCR amplification duplicates were removed via Samtools (Version 0.1.19-44428cd). Freebayes (Version 0.9.14-19-g8a407cf) was employed for variant calling in the hexaploid genome.

For haplotyping, our method takes FASTA, SAM and VCF formatted files as inputs. We base the reconstruction on seed regions, which are small sets of polymorphic sites. For example, 3 polymorphic sites with 2 alleles each, would allow for up to 2x2x2=8 haplotypes. Only a subset of those will be supported by reads. The algorithm searches for all possible seed regions containing six or more sequence patterns. Seeds, however, can be interleaved. Haplotypes in a seed region are sorted by the number of their supporting reads. Different seed regions are sorted according to the number of supporting reads for the sixth-most strongly supported haplotype, because we expect 6 haplotypes.

The following steps are done iteratively on each seed region from the sorted seed list. The six most supported sequence patterns in each seed region are considered as the six haplotype cores. Then, each haplotype is built up on each core according to the supporting reads for that sequence pattern (Figure 3). The haplotypes are extended via uniquely matched reads. A uniquely matched read is a read that matches exactly one haplotype while being distinct from the other haplotypes in the seed region. Chaining haplotypes through uniquely matched reads produces six haplotypes in an extended seed region (Supplementary Figure 8a and b).

Then the adjacent overlapping seed regions are merged (Supplementary Figure 8c). One haplotype in a seed region is merged to a haplotype in the adjacent seed region if they share a uniquely matched overlapping sequence with each other. Thereafter, we utilize paired-end reads to connect the haplotypes obtained so far. All the Illumina raw reads are mapped back to the phased haplotypes. Only perfectly matched paired-end reads are considered as haplotype connections. The inner-scaffold haplotyping is done exploiting the paired-end reads within one scaffold. The algorithm starts with the highest supported connection and merges the connected haplotypes as a new haplotype. Possible conflicts are checked for and avoided in this step. A conflict would occur when two haplotypes in one seed region are connected via a path through the haplotypes in other seed regions. Paired-end reads are further utilized for inter-scaffold connection to elongate and merge the haplotypes from different scaffolds. The haplotyping method utilizes parallel computation in order to speed up the analysis.

**Supplementary Figure 8:**
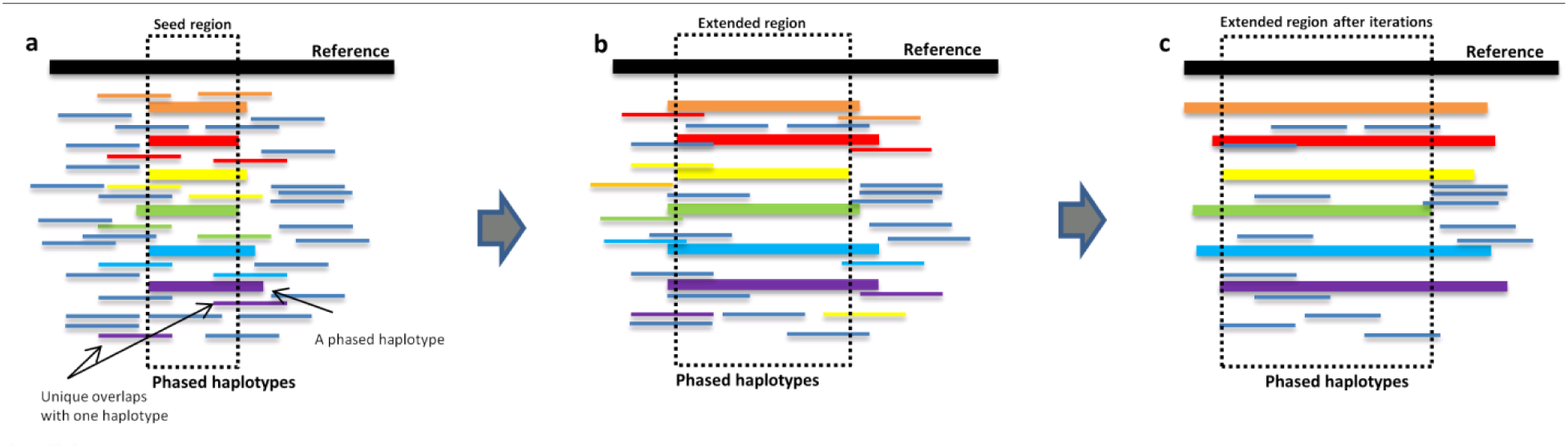
The seed formation and seed extension process. **(a)** A constructed seed (dashed box) and its corresponding haplotypes (different colors). (b) Seed extension with the reads that are uniquely mapped with the constructed haplotypes. (c) A region with 6 constructed haplotypes, they are obtained after the iteration of seed extension.

### Haplotype evaluation using 454 reads

To evaluate the haplotyping accuracy, we have used a set of 454 (Supplementary Table 1, A454) reads that had been produced earlier but was not utilized for assembly or phasing. Each 454 read can be considered as a short DNA fragment from one chromosome, except for some chimeric reads in rare cases. Roche 454-trimmed reads were mapped against HI-assembly. Only the polymorphic sites indicated by variant calling were extracted and their overlaps with haplotypes were evaluated. The “Match” and “Mismatch” sites of each overlap were recorded for further analysis and visualization.

### HI-assembly and variant correction

A python script is employed to convert all inter-scaffolds connection information into SSPACE tab format. Exhaustive comparisons among scaffolds using blast were employed to identify and remove redundancy in scaffolds. If one scaffold is already covered by another longer scaffold with more than 85% sequence identity and more than 85% sequence length, the shorter one was removed. Additional manual checking of long candidates (>10kb) via Circos visualization was also included. Finally, the non-redundant scaffold sequences without the endophyte *Bacillus pumilus* genome, were employed as input scaffolds to SSPACE (Version 3.0) for scaffolding. The library settings for SSPACE scaffolding were based on the insert size distribution of all paired-end libraries (Supplementary Figure 2).

We refined variants according to phased haplotypes. There are three possibilities for one phased variant: (1) all six haplotypes covered the variant, (2) some of the haplotypes covered the variant, and (3) the variant was located outside of haplotype blocks. There is insufficient information to update alleles in the second and third categories. Our method distinguishes the first group from the other two and categorizes its members in the following subgroups: (a) all alleles are the same and (b) the alleles are different. The first subgroup is put aside as an error in variant calling, which means that there is insufficient support to consider this position as a variant position. We therefore removed the variant calling result of this position from the original VCF file. For the second group, our method ranks alleles firstly based on the number of supporting haplotypes, secondly according to the number of supporting reads. Then the first allele is picked as a reference allele and the rest are considered as alternative alleles.

### Genome annotation

Six transcriptome data sets from four previous studies (Wang et al. 2010; Tao et al. 2012; Xie et al. 2012; Firon et al. 2013) and two additional RNA-seq experiments were mapped on scaffolds using HISAT (Version 0.1.5-beta, Kim et al. 2015). These transcriptome data represented different plant tissues such as leaves, petioles, stems, and roots from different development stages. All gene models were extracted by StringTie (Pertea et al. 2015). The obtained transcripts were further annotated using homologous protein searching in public database Tair and Uniprot.

NCBI proteins from *Nicotiana sylvestris*, *Nicotiana tomentosiformis*, *Solanum indicum*, *Solanum tuberosum*, *Solanum lycopersicum*, *Nicotiana tabacum* and *Sesamum indicum* genomes were mapped against the present genome of *Ipomoea batatas*, using SPALN (Iwata and Gotoh 2012) (Version 2.2.0). All aligned protein locations were considered as potential genes in HI-assembly. RepeatMasker (Version open-4.0.5) was used to mask and classify repeat sequences in the genome. RepeatModeler (Version 1.0.8, Smit and Hubley) was employed to identify novel repeat sequences.

### Phylogenetic analysis

A python script was employed to extract alignments from all phased regions. MEGA-Computing Core (Kumar et al. 2012) was utilized to compute all the phylogenetic trees of extracted alignments on computer farm. Trees were classified into groups based on their topology structures and average haplotype lengths in these groups were compared. The dominant tree structure was selected and a consensus tree with average branch lengths was obtained.

### Data accessibility

The *Ipomoea batatas* genome sequence, including consensus scaffolds, haplotype-resolved scaffolds, and unplaced contigs, are publicly available at the *Ipomoea batatas* genome browser http://public-genomes-ngs.molgen.mpg.de/SweetPotato/. Two transcriptome data sets and one sequenced cDNA library are also available for download. cDNA clone delivery is also possible upon request. The HI-assembly and the WGS raw data have been deposited with European Nucleotide Archive (ENA) under project number PRJEB14638 and National Center for Biotechnology Information (NCBI) under project number PRJNA301667.

## Acknowledgments

We thank Dr. Junyi Dai and Dr. Zoran Nikoloski for helpful discussions. Dr. Jun Yang acknowledges supports from the Alexander von Humboldt Foundation (Forschungsstipendium für erfahrene Wissenschaftler). Mr. M-Hossein Moeinzadeh acknowledges supports from IMPRS-CBSC dortoral program. This project was funded by the National Natural Science Foundation of China (31201254, 31361140366) the International Science & Technology Cooperation Program of China (2015DFG32370), the National High Technology Research and Development Program of China (2011AA100607-4, 2012AA101204-3), the Chinese Academy of Sciences (2012KIP518), the China Postdoctoral Science Foundation (2012M520945), the Shanghai Municipal Afforestation & City Appearance and Environmental Sanitation Administration (G102410, F122422, F132427, G142434, G152429), and the Science and Technology Commission of Shanghai Municipality (14DZ2260400).

## Author contributions

J. Y., M-H. M., H. K., A. R. F., B. T., M. V. and P. Z. planned and coordinated the project and wrote the manuscript. G. -L. L., J. -L. Z. and Z. S. supplied the newly bred cultivar, Taizhong6. W. -J. F., G. -F. D. and H. -X. W. prepared genomic DNA. H. K. conducted the primary genome assembly and repeat sequence identification. J. Y. and M-H. M. conducted haplotyping and genome evolution analysis. J. H., P. X. and F. -H. H. supported and inspired a part of the analysis.

## Competing financial interests

The authors declare no competing financial interests.

## Reference

Aguiar D, Istrail S. 2013. Haplotype assembly in polyploid genomes and identical by descent shared tracts. Bioinformatics 29(13): 352–360.

Berger E, Yorukoglu D, Peng J, Berger B. 2014. Haptree: A novel bayesian framework for single individual polyplotyping using NGS data. PLOS Computational Biology 10(3): e1003502. doi: 10.1371/journal.pcbi.1003502

Boetzer M, Henkel CV, Jansen HJ, Butler D, Pirovano W. 2011. Scaffolding pre-assembled contigs using SSPACE. Bioinformatics 27(4): 578–579.

Boycheva S, Daviet L, Wolfender J-L, Fitzpatrick TB. 2014. The rise of operon-like gene clusters in plants. Trends in plant science 19(7): 447–459.

Cao H, Wu H, Luo R, Huang S, Sun Y, Tong X, Xie Y, Liu B, Yang H, Zheng H. 2015. De novo assembly of a haplotype-resolved human genome. Nature biotechnology 33(6): 617–622.

Choulet F, Alberti A, Theil S, Glover N, Barbe V, Daron J, Pingault L, Sourdille P, Couloux A, Paux E. 2014. Structural and functional partitioning of bread wheat chromosome 3B. Science 345(6194). doi: 10.1126/science.1249721.

Consortium IH. 2005. A haplotype map of the human genome. Nature 437(7063): 1299–1320.

Consortium IWGS. 2014. A chromosome-based draft sequence of the hexaploid bread wheat (Triticum aestivum) genome. Science 345(6194). doi: 10.1126/science.1251788.

Consortium PGS. 2011. Genome sequence and analysis of the tuber crop potato. Nature 475(7355): 189–195.

Duitama J, McEwen GK, Huebsch T, Palczewski S, Schulz S, Verstrepen K, Suk E-K, Hoehe MR. 2011. Fosmid-based whole genome haplotyping of a HapMap trio child: evaluation of Single Individual Haplotyping techniques. Nucleic acids research. doi: 10.1093/nar/gkr1042.

Fernie AR, Tohge T. 2015. Location, location, location–no more! The unravelling of chromatin remodeling regulatory aspects of plant metabolic gene clusters. New Phytologist 205(2): 458–460.

Firon N, LaBonte D, Villordon A, Kfir Y, Solis J, Lapis E, Perlman TS, Doron-Faigenboim A, Hetzroni A, Althan L. 2013. Transcriptional profiling of sweetpotato (Ipomoea batatas) roots indicates down-regulation of lignin biosynthesis and up-regulation of starch biosynthesis at an early stage of storage root formation. BMC genomics 14(1). doi: 10.1186/1471-2164-14-460.

Garrison E, Marth G. 2012. Haplotype-based variant detection from short-read sequencing. arXiv preprint. arXiv: 10.1093/nar/gks708.

Hirakawa H, Okada Y, Tabuchi H, Shirasawa K, Watanabe A, Tsuruoka H, Minami C, Nakayama S, Sasamoto S, Kohara M. 2015. Survey of genome sequences in a wild sweet potato, *Ipomoea trifida (HBK) G*. Don. DNA Research. doi: 10.1093/dnares/dsv002.

Iwata H, Gotoh O. 2012. Benchmarking spliced alignment programs including Spaln2, an extended version of Spaln that incorporates additional species-specific features. Nucleic acids research. doi: 10.1093/nar/gks708.

Jia J, Zhao S, Kong X, Li Y, Zhao G, He W, Appels R, Pfeifer M, Tao Y, Zhang X. 2013. Aegilops tauschii draft genome sequence reveals a gene repertoire for wheat adaptation. Nature 496(7443): 91–95.

Kajitani R, Toshimoto K, Noguchi H, Toyoda A, Ogura Y, Okuno M, Yabana M, Harada M, Nagayasu E, Maruyama H. 2014. Efficient de novo assembly of highly heterozygous genomes from whole-genome shotgun short reads. Genome research 24(8): 1384–1395.

Kim D, Langmead B, Salzberg SL. 2015. HISAT: a fast spliced aligner with low memory requirements. Nature methods 12(4): 357–360.

Kim S-H, Hamada T. 2005. Rapid and reliable method of extracting DNA and RNA from sweetpotato, *Ipomoea batatas* (L). Lam. Biotechnology letters 27(23–24): 1841–1845.

Kriegner A, Cervantes JC, Burg K, Mwanga RO, Zhang D. 2003. A genetic linkage map of sweetpotato [*Ipomoea batatas* (L.) Lam.] based on AFLP markers. Molecular Breeding 11(3): 169–185.

Kumar S, Stecher G, Peterson D, Tamura K. 2012. MEGA-CC: computing core of molecular evolutionary genetics analysis program for automated and iterative data analysis. Bioinformatics 28(20): 2685–2686.

Kyndt T, Quispe D, Zhai H, Jarret R, Ghislain M, Liu Q, Gheysen G, Kreuze JF. 2015. The genome of cultivated sweet potato contains Agrobacterium T-DNAs with expressed genes: An example of a naturally transgenic food crop. Proceedings of the National Academy of Sciences 112(18): 5844–5849.

Li F, Fan G, Lu C, Xiao G, Zou C, Kohel RJ, Ma Z, Shang H, Ma X, Wu J. 2015. Genome sequence of cultivated Upland cotton (Gossypium hirsutum TM-1) provides insights into genome evolution. Nature biotechnology 33(5): 524–530.

Li F, Fan G, Wang K, Sun F, Yuan Y, Song G, Li Q, Ma Z, Lu C, Zou C. 2014. Genome sequence of the cultivated cotton Gossypium arboreum. Nature genetics 46(6): 567–572.

Ling H-Q, Zhao S, Liu D, Wang J, Sun H, Zhang C, Fan H, Li D, Dong L, Tao Y. 2013. Draft genome of the wheat A-genome progenitor Triticum urartu. Nature 496(7443): 87–90.

Luo R, Liu B, Xie Y, Li Z, Huang W, Yuan J, He G, Chen Y, Pan Q, Liu Y. 2012. SOAPdenovo2: an empirically improved memory-efficient short-read de novo assembler. GigaScience 1(1). doi:10.1186/2047-217X-1-18

Margulies M, Egholm M, Altman WE, Attiya S, Bader JS, Bemben LA, Berka J, Braverman MS, Chen Y-J, Chen Z. 2005. Genome sequencing in microfabricated high-density picolitre reactors. Nature 437(7057): 376–380.

Nützmann HW, Osbourn A 2015. Regulation of metabolic gene clusters in Arabidopsis thaliana. New Phytologist 205(2): 503–510.

Ossowski S, Schneeberger K, Lucas-Lledó JI, Warthmann N, Clark RM, Shaw RG, Weigel D, Lynch M. 2010. The rate and molecular spectrum of spontaneous mutations in Arabidopsis thaliana. Science 327(5961): 92–94.

Ozias-Akins P, Jarret RL. 1994. Nuclear DNA content and ploidy levels in the genus *Ipomoea*. Journal of the American Society for Horticultural Science 119(1): 110–115.

Peng Y, Leung HC, Yiu S-M, Chin FY. 2012. IDBA-UD: a de novo assembler for single-cell and metagenomic sequencing data with highly uneven depth. Bioinformatics 28(11): 1420–1428.

Pertea M, Pertea GM, Antonescu CM, Chang T-C, Mendell JT, Salzberg SL. 2015. StringTie enables improved reconstruction of a transcriptome from RNA-seq reads. Nature biotechnology 33(3): 290–295.

Schafleitner R, Tincopa LR, Palomino O, Rossel G, Robles RF, Alagon R, Rivera C, Quispe C, Rojas L, Pacheco JA. 2010. A sweetpotato gene index established by de novo assembly of pyrosequencing and Sanger sequences and mining for gene-based microsatellite markers. BMC genomics 11(1). doi: 10.1186/1471-2164-11-604

Slater GS, Birney E 2005. Automated generation of heuristics for biological sequence comparison. BMC bioinformatics 6(1). doi: 10.1186/1471-2105-6-31

Smit A, Hubley R. RepeatModeler (2008–2010) RepeatModeler-1.0.8 Available: http://www.repeatmasker.org/RepeatModeler.html. Accessed 2015 October.

Snyder MW, Adey A, Kitzman JO, Shendure J. 2015. Haplotype-resolved genome sequencing: experimental methods and applications. Nature Reviews Genetics 16(6): 344–358.

Tao X, Gu Y-H, Wang H-Y, Zheng W, Li X, Zhao C-W, Zhang Y-Z. 2012. Digital gene expression analysis based on integrated de novo transcriptome assembly of sweet potato [Ipomoea batatas (L.) Lam.]. PloS one 7(4). doi: 10.1371/journal.pone.0036234.

Ukoskit K, Thompson PG. 1997. Autopolyploidy versus allopolyploidy and low-density randomly amplified polymorphic DNA linkage maps of sweetpotato. Journal of the American Society for Horticultural Science 122(6): 822–828.

Wang K, Wang Z, Li F, Ye W, Wang J, Song G, Yue Z, Cong L, Shang H, Zhu S. 2012. The draft genome of a diploid cotton Gossypium raimondii. Nature genetics 44(10): 1098–1103.

Wang Z, Fang B, Chen J, Zhang X, Luo Z, Huang L, Chen X, Li Y. 2010. De novo assembly and characterization of root transcriptome using Illumina paired-end sequencing and development of cSSR markers in sweetpotato (Ipomoea batatas). BMC genomics 11(1): 726. doi: 10.1186/1471-2164-11-726.

Xie F, Burklew CE, Yang Y, Liu M, Xiao P, Zhang B, Qiu D. 2012. De novo sequencing and a comprehensive analysis of purple sweet potato (*Ipomoea batatas* L.) transcriptome. Planta 236(1): 101–113.

Zimin AV, Marçais G, Puiu D, Roberts M, Salzberg SL, Yorke JA. 2013. The MaSuRCA genome assembler. Bioinformatics 29(21): 2669–2677.

